# JNJ-42153605, a mGluR2 PAM, potentiates Levetiracetam treatments of TBI to mitigate subsequent tau aggregation in a larval zebrafish model

**DOI:** 10.64898/2026.06.20.733541

**Authors:** Laszlo F. Locskai, Shayleen Ghassemi, Samantha A.W. Tan, Melissa J. Kinley, W. Ted Allison

**Affiliations:** Department of Biological Sciences, University of Alberta, Edmonton AB, T6G 2E9, Canada; Centre for Prions & Protein Folding Disease, University of Alberta, Edmonton AB, T6G 2M8, Canada

**Keywords:** anti-convulsant drugs, neurotrauma, neurodegeneration, cognitive decline, neural hyperexcitability, synaptic vesicle protein 2a, mGlu2, post-traumatic epilepsy, tauopathy

## Abstract

Traumatic brain injury (TBI) has long-term consequences that include chronic traumatic encephalopathy (CTE) and an elevated risk for Alzheimer Disease (AD). These dementias ultimately manifest as tauopathies but may begin with acute neuronal dysfunction including post-traumatic seizures. Provocative evidence suggests that these prodromal seizures are a viable target to mitigate the later onset of dementias, and anti-epileptic drugs (AED) that increase the threshold of action potentials have indeed been shown to mitigate later tauopathies[1, 2]. Here, we test whether AEDs and other compounds that modulate synaptic transmission, applied immediately after TBI, can also act as prophylactics that acutely block subsequent CTE-like tau aggregation and neurodegeneration in a larval zebrafish model expressing a Tau-GFP fusion protein in the CNS. Levetiracetam (LEV) is an AED that modulates synaptic vesicle release. Application of LEV immediately following TBI abrogated TBI-induced tau aggregation (IC_50_ = 3.168×10^-3^ mM) and cell death in the larval zebrafish TBI model. We next considered a polypharmacy approach involving metabotropic glutamate receptor 2 (mGluR2), because mGluR2 positively allosteric modulators (PAMs) such as JNJ-42153605 have previously been able to improve LEV’s action in reducing some recalcitrant forms of seizure in a mouse model. We found that JNJ-42153605 was itself effective at blocking TBI-induced tau aggregation (IC50 = 8.691 ×10^-5^ mM). Moreover, a subeffective dose of JNJ-42153605 (10^-5^ mM) was able to substantially improve the efficacy of LEV (∼16-fold) in its prophylactic actions. Thus, LEV and JNJ-42153605 applied briefly after TBI offer a potent polypharmacy approach, at least in our preclinical animal model, to tackle the later tau aggregation and neurodegeneration that follows from TBI neurotrauma. These results warrant further investigation, including testing in mammalian TBI models with longer disease course and more conventional markers of tau pathology.

**Highlights:**

- A larval zebrafish model of blast TBI provides good sample sizes and throughput
- Expressing Tau fused to GFP in zebrafish enables in vivo quantification
- Levetiracetam applied following TBI reduced cell death and tau aggregation
- An mGluR2 PAM applied after TBI potentiated the benefits of Levetiracetam
- Anti-epileptics targetted at synapse physiology have promise in this paradigm

## INTRODUCTION

Dementias such as AD and CTE progress inexorably from behavioural and psychological symptoms of dementia towards death. Among other pathologies, these dementia outcomes are the manifestation of progressive tauopathy burden that are characterized by the templated misfolding and accumulation of aggregated forms of the tau protein. Treatments that can slow the progression of dementia are rare. In part this is because early diagnosis of dementia is challenging, yet a growing consensus agrees that it is essential to intervene early in the disease course. Intriguingly, several prominent dementias share head injury as a cause or leading risk factor. This suggests that interventions immediately following TBI may be able to disrupt the earliest pathomechanisms in various dementias, and thereby prevent or slow the onset of symptoms[2].

Acute events following TBI prominently include post-traumatic seizures (PTS). Emerging evidence indicates PTS are prodromal of later dementias[2]. Reducing seizure-like events and other neural hyperexcitability following TBI by short-term AED application is sufficient to prevent later tau aggregation and cell death in zebrafish models[1]. Similarly, briefly applying AED following blast TBI in mouse models was sufficient to reduce cognitive decline and sleep disruption many months after neurotrauma[3]. Moreover, increasing seizures following TBI results in increased tauopathy and can reverse the AED action[1]. Importantly then, AEDs applied following TBI offer hope to prophylactically prevent later dementias[2]. Here we consider what alternative classes and regimens of AED have the greatest potential to mitigate later tauopathies.

Strategies to disrupt the early-onset pathomechanisms of dementia have often revolved around synaptic dystrophies, because synapse disruption is one of the earliest observable events presaging cognitive decline. To date, the principal AED tested as a dementia prophylactic is retigabine[1, 3], which instead acts in the axon initial segment to reduce seizures by increasing the threshold of action potential firing, via altering voltage-gated potassium K_v_7 channel properties. Here, we test the prophylactic potential of AEDs that operate in the presynaptic compartment and reduce synaptic transmission. Several such AEDs are applied clinically to reduce PTS, offering good potential for clinical translation and for repurposing of AEDs into dementia prophylactics.

A synthesis of available literature led us to study levetiracetam (LEV) as a potential dementia prophylactic[2]. LEV binds to synaptic vesicle protein 2a (SV2a), reducing synaptic vesicle release, and this is thought to be its primary mode of action reducing seizure outcomes. Several lines of evidence suggest the LEV has additional mechanisms of action, including that animals lacking SV2a exhibit greatly reduced, but not completely absent, responses to LEV application[4]. LEV is effective in mitigating PTS[5–13]. Intriguingly, LEV has also shown early promise to slow the progression of dementias in subsets of mild cognitive impairment (MCI) patients demonstrating seizure-like activity [2, 14], though its broad applicability across dementias is not yet established[15].

Here, we demonstrate that LEV, applied briefly following TBI, has good potential to prevent later cell death and tau aggregation, at least in a preclinical animal model. LEV treatment is not without its drawbacks, and so we sought additional AED treatments that might potentiate LEV to mitigate cell death and tau aggregation following TBI. Conceptually, a polypharmacy approach following TBI could preserve LEV’s efficacy at lower doses, reducing potential side-effects or the development of drug tolerance/insensitivity. We focus on pharmacology against metabotropic glutamate receptor 2 (mGluR2) as a presynaptic target that modulates synaptic vesicle release. Elegant studies with a series of mGluR2 PAMs show that they act synergistically with LEV to prevent seizures in a therapy-resistant model, the 6 Hz mouse model of psychomotor seizures[16, 17]. Our rationale extends directly from those past works, and to aims to explore potentiated signaling at the intersection of two known presynaptic pathways that modulate glutamate release, SV2a and mGluR2. Moreover, mGluR2 PAMs do not directly activate the receptor at the ligand binding site and this may provide advantages, including a minimized alteration to essential glutamate signalling and reduced tachyphylaxis[16, 17]. Here, we find that an mGluR2 PAM, JNJ-42153605, is itself able to prevent later tau aggregation, and potentiate the effects of LEV, when applied briefly after TBI.

## RESULTS

### Levetiracetam applied after TBI prophylactically prevents subsequent tau aggregation

The events following TBI neurotrauma involve various forms of neural hyperexcitability including post-traumatic seizures, and mitigating this neural hyperexcitability is a promising route to mitigate the later onset of tauopathies[2]. LEV is an anti-epileptic drug (AED) with some reported promise in mitigating similar dementia outcomes[14, 15, 18]; we sought to appreciate if LEV can lessen the tau aggregation burden that is induced by TBI. We turned to larval zebrafish as a previously published preclinical model of TBI and tau aggregation, because this model enables robust sample sizes and has previously proven able to identify AED candidates that mitigate TBI outcomes, and those AEDs translate well into rodent models for blocking tau aggregation development[1–3].

We quantified tau aggregates in each individual animal as the abundance of Tau4R-GFP puncta in the CNS; this “tau reporter” line of fish is transparent and expresses the aggregation-prone 4R portion of the human Tau protein fused to green fluorescent protein (GFP) in neurons (**Fig. 1A**) and has been thoroughly validated as a faithful reporter of tau aggregation [1,19,20]. We quantified tau puncta in the spinal cord as they are reliably visible in this thinner portion of the CNS, allowing less preparation per animal and relatively large sample sizes. The abundance of tau puncta in the spinal cord is a reliable proxy for tau burden in the brain, insomuch that these two metrics correlate well with each other – they both increase with various doses of injury and with time following injury[1]. TBI was delivered as a pressure wave through the larval zebrafish’s bath media[1, 19, 20], in a format most analogous to blast pressure waves moving through all organs that occurs during TBI in patients experiencing neurotrauma following their proximity to explosive blasts.

**Figure 1.**
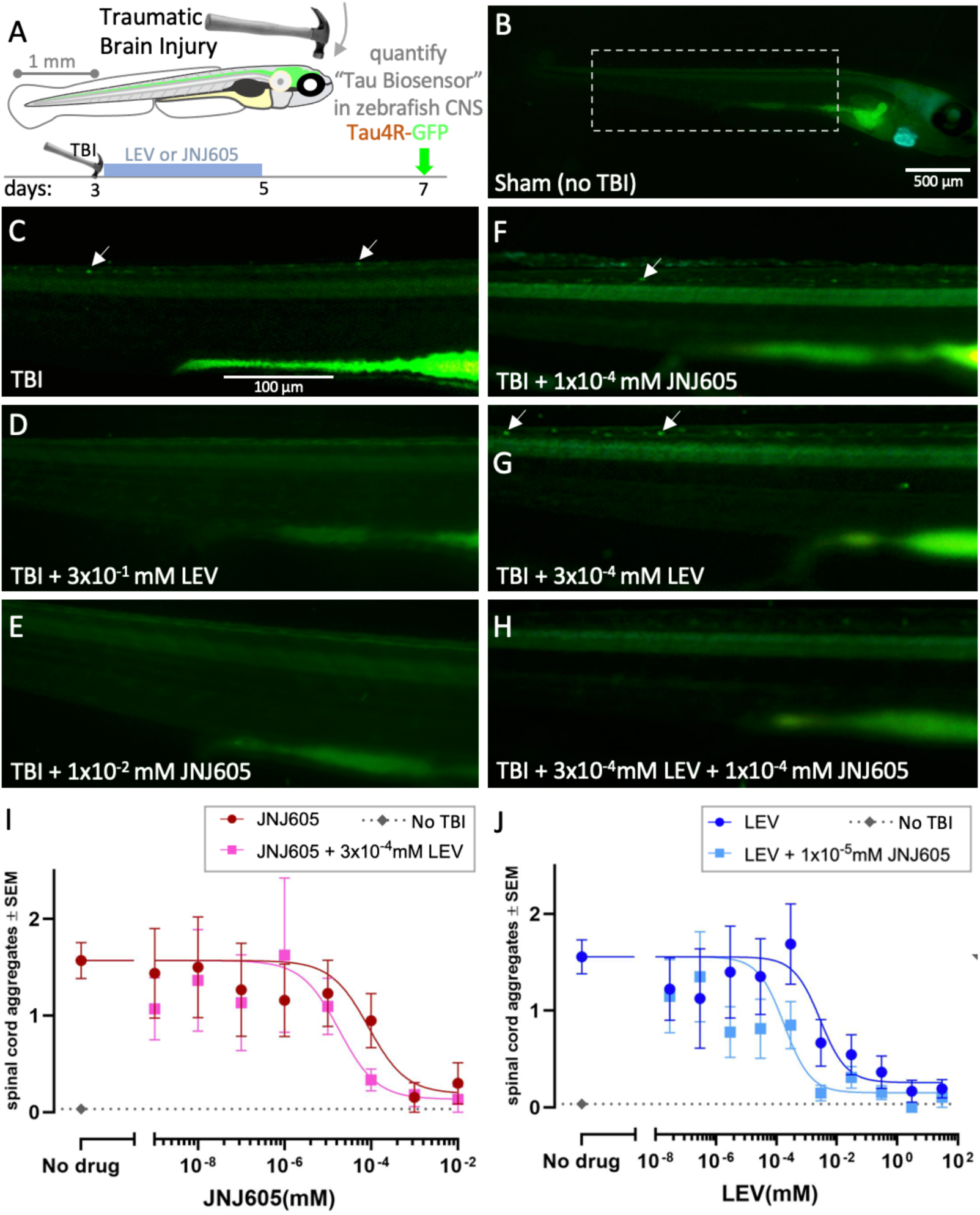
Levetiracetam and an mGluR2 PAM, JNJ-42153605, potentiate each other’s ability to prevent subsequent tau aggregation when applied following TBI. Tau4R-GFP in the CNS neurons of zebrafish larvae receiving traumatic brain injury (TBI) report increased tau puncta (A, C), examples indicated by arrows. At high doses anti-epileptic drug Levetiracetam (LEV) or JNJ-42153605 (“JNJ605”, a mGluR2 positive allosteric modulator [PAM]) applied immediately following TBI substantially reduced tau aggregation to the levels of uninjured animals (D, E, I, J). Lower doses of LEV or JNJ605 were less effective but potentiated each other to reduce tau aggregation to baseline levels of uninjured animals (F-J). Scale bar in panel C applies for panels D-H. I. Treatment with JNJ605 following TBI significantly reduced subsequent tau aggregation (IC50 8.691 ×10^-5^ mM, 95% CI 2.086-28.06 ×10^-5^ mM, Nagelkerke’s pseudo-R-squared 0.07615) and a sub-effective dose of LEV (3×10^-4^ mM) improved the efficacy of JNJ605 about 5-fold (3.3-fold to 5.8-fold using 95% CI) (1.816×10^-5^ mM, 95% CI 6.361-48.80 ×10^-6^ mM; p=0.0455, pseudo-R-squared 0.1411), with each curve being significantly different (p=0.0455). J. In trials that were independent from those in panel I, treatment with LEV following TBI significantly reduced subsequent tau aggregation (IC50 3.168 ×10^-3^ mM, 95% CI 7.378-187.7 ×10^-4^ mM, pseudo-R-squared 0.1836). A subeffective dose of JNJ605 (10^-5^ mM) improved the efficacy of LEV by about 10-fold or more (14 to 31 fold using 95% CI) (IC50 of 1.981 ×10^-4^ mM, 95% CI 5.447-61.31 ×10^-5^ mM, pseudo-R-squared 0.1982), with each curve being significantly different (p=0.0003). n>8 animals per dose as detailed with the raw data plotted in Supplemental Figure S1. A separate analysis of the same data is in Fig S2. Dose response curves were generated using three parameter inhibitor vs response nonlinear regression. Statistical differences between dose response curves were assessed using Nagelkerke’s likelihood ratio test.

Following delivery of TBI at 3 days post-fertilization (dpf), we observed increased tau puncta at 7dpf compared to sham treated animals (**Fig 1B,C**. see also raw data plotted in **Supp Fig S1**), as expected[1]. To test the actions of LEV we applied it for 38-44 hours, beginning within 30-90 minutes following TBI, by adding it to the bathing media that houses the larvae. This application route, adding AEDs to the bath media of larval zebrafish, matches past (high-throughput) efforts to dissect acute seizures in zebrafish, although it is noteworthy that our approach utilizes an extended duration of AED application.

LEV prevented the development of tau puncta induced by TBI in a dose-dependent manner (**Fig 1D,G,J dark blue line**; IC50 3.168 ×10^-3^ mM, 95% confidence interval (CI) 7.378-187.7 ×10^-4^ mM. Raw data plotted in **Supp Fig S1C**). An alternative analysis of the same data, considering what percentage of animals had tau aggregation considerably above baseline levels (>1 tau puncta per individual) supported these findings showing that LEV increased the odds of larvae not having detectable puncta post-TBI 1.11-fold/mM (**Supp Fig S2B.** odds ratio (OR) 1.11, 95% CI 1.041-1.271).

### mGluR2 PAM JNJ605 potentiates the action of LEV in this model

We investigated the potential potentiation between the two compounds by applying LEV together with a subeffective dose of JNJ605 (10^-5^ mM, a dose with no overt impact in our assay, Figure 1I, Supp Fig S1A, S2A) and quantifying tau aggregates induced by TBI. First, we established the dose relationship to tau aggregation with LEV and with JNJ605, showing LEV to be effective at concentrations greater than 3×10^-4^ (**Fig 1D,G,J dark blue line**; Raw data plotted in **Supp Fig S1C**), and JNJ to be effective at doses greater than 1×10^-4^ (**Fig. 1E,F,I dark red line**. Raw data in **Supp Fig S1A**). We then determined the ability of JNJ605 to potentiate LEV, and vice versa in a separate experiment with different animals. In the presence of a fixed concentration at a sub-effective dose of LEV (10^-4^ mM), we observed a reduction in punctae at concentrations of JNJ605 as low as 10^-8^ (**Figure 1 H,I magenta line**). Similarly, in the context of a fixed, but sub effective, concentration of JNJ605 of 10^-5^ mM we observed efficacy of LEV down to 10^-8^ mM (**Fig1 G,H,J** compare light and dark blue lines in panel J). Likelihood ratio tests comparing if dose response curves were significantly different, P = 0.0003. Raw data is in **Supp Fig S1D**). Re-analyzing the same animals in Figure 1I further supports this conclusion by showing that the proportion of animals exhibiting tau aggregation was reduced by LEV in a dose-dependent fashion, and the efficacy of LEV was improved by the presence of JNJ605 which increased the odds of larvae being tau free post-TBI 2.7-fold (**Supp Fig S2B.** OR 2.694, 95% CI 1.674-4.459). Re-analysis of the same animals in Figure 1J provided an equivocal interpretation where LEV provided an increase in JNJ605 efficacy that was non-significant (**Supp Fig S2A**. OR 1.443, 95% CI 0.8702-2.443); one of several possibilities is that the sub-effective dose of LEV selected was not optimal for consistently improving the impacts of JNJ605.

The 11-point dose-response curve attained for LEV in the presence of JNJ605 was not distinctly unimodal (**Fig 1J**, light blue squares). In the presence of JNJ605, LEV doses in the range of 10^-5^ to 10^-3^ resulted in substantially lower tau aggregation compared to the same doses of LEV alone, whereas the tau aggregation levels were less impacted by JNJ605 at lower doses of LEV. Regardless, data compiled from multiple experiments assessing hundreds of individuals in total, demonstrate that several low doses of LEV (10^-5^ to 10^-3^ mM) were substantially more successful in blocking tau aggregation when in the presence of subeffective dose of JNJ605 (**Fig 1J**) and conservatively suggest a >10-fold improvement in LEV efficacy (between 14-fold and 31-fold change is estimated from the CI). In sum, the data support a strong potential for an mGluR2 PAM, JNJ605, to improve the actions of LEV towards mitigating tau aggregation following neurotrauma in our model. Independent experiments confirmed this property of JNJ605 regardless of its source of synthesis (**Supp Figs S3 and S4**).

### JNJ605 and LEV reduce CNS cell death following TBI

We next determined if LEV and JNJ605 are able to reduce neurotrauma-induced cell death in the CNS. We applied the same TBI paradigm as above but with a new endpoint that quantified cell death via immunohistochemistry against activated-caspase3 at two days post-injury. TBI led to approximately a three-fold increase in the level of CNS cell death (p<0.0001) compared to uninjured control animals (**Fig 2**), as expected[1]. Akin to previous AEDs applied immediately after TBI[1], both LEV and JNJ605 were able to reduce subsequent CNS cell death. Higher doses of each compound(3×10^-1^ and 10^-3^ mM, respectively) resulted in TBI-treated larvae exhibiting cell death at the same abundance as uninjured sham animals (**Fig 2E**). Lower doses of LEV and JNJ605 (3×10^-4^ and 10^-5^ mM, respectively) were moderately effective (**Fig 2F**). Combining these lower doses of LEV and JNJ605 resulted in TBI-induced cell death being reduced by ∼3-fold (p<0.0001), to levels akin to uninjured sham animals (**Fig 2F**). Overall, LEV and the mGluR2 PAM JNJ605 appear able to potentiate each other’s protective effects on CNS cell survival, akin to their impact on tau puncta above.

**Figure 2.**
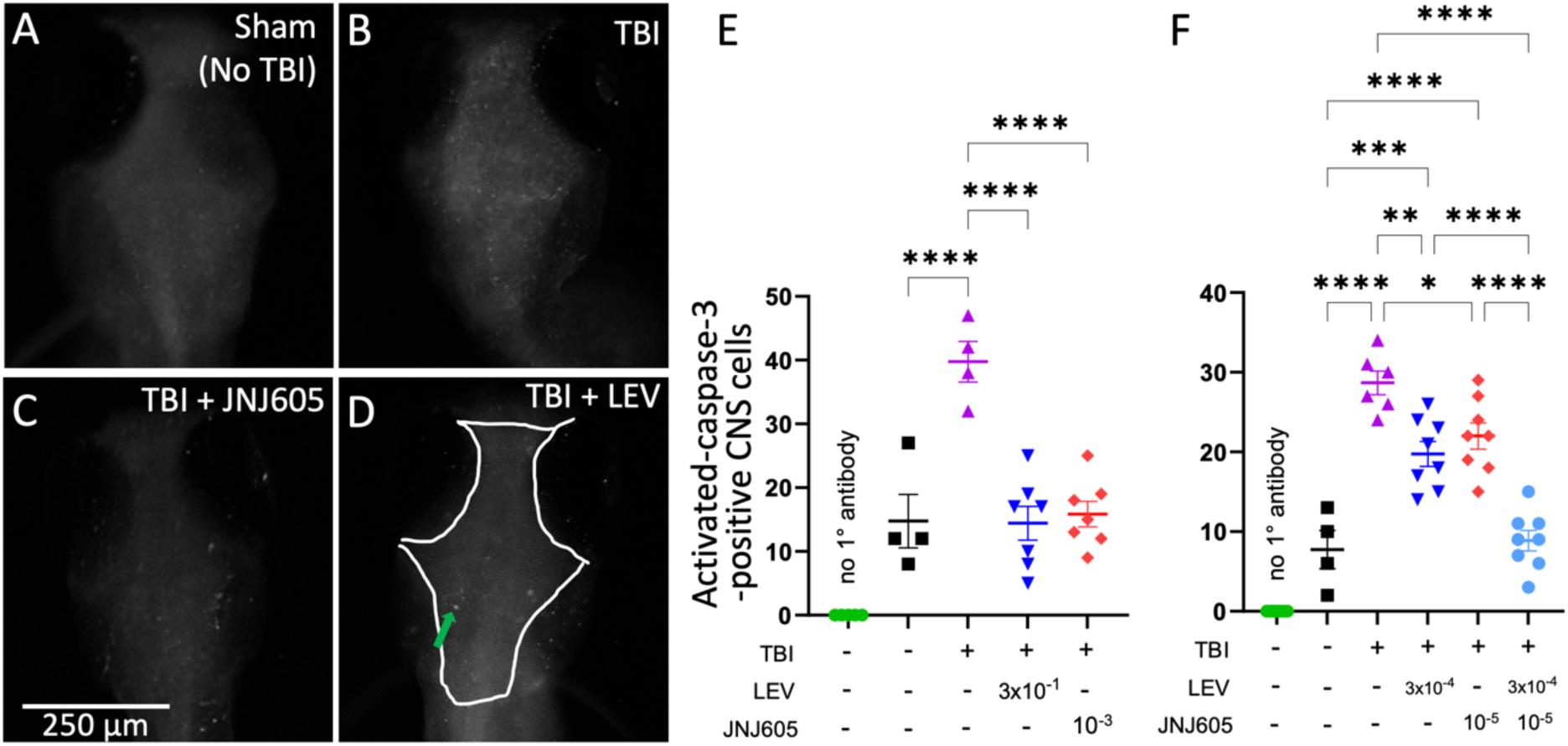
Levetiracetam and the mGluR2 PAM JNJ-42153605 interact to mitigate TBI-induced CNS cell death. Traumatic brain injury (TBI) delivered to larval zebrafish induces cell death several days later (detected by anti-activated-Caspase3; compare panel A vs. B, dorsal view of larval zebrafish brains). Higher doses of Levetiracetam (LEV, 3×10^-1^ mM) or JNJ-42153605 (“JNJ605” at 10^-3^ mM) delivered for 44 hours post-injury reduced cell death to baseline levels (C-E). Lower doses of LEV or JNJ605 were less effective individually, but when combined LEV and JNJ605 mitigated the impacts of TBI on subsequent CNS cell death. Each symbol represents an individual animal, mean ±SE plotted. *p<0.05, **p<0.01, ***P<0.001, ****p<0.0001 by one-way ANOVA and Tukey’s multiple comparisons test post-hoc.

### mGluR2 PAM specificity of action

We next probed the specificity of JNJ605 in our paradigm, testing other mGluR2 PAMs, compounds inactive as mGluR2 PAM, and mGluR2 negative allosteric modulators (NAMs).

We determined that the mGluR2 PAM JNJ-46356479 (“JNJ479”)[16, 21] is able to reduce TBI-induced cell death back to baseline levels (**Fig 3A**). JNJ479 and JNJ605 have distinct backbone structures (**Fig 3D**), suggesting it is unlikely that they would share any imagined off-target effects. Instead, it is likely they both act principally through a shared allosteric modulation of the mGluR2 protein.

**Figure 3.**
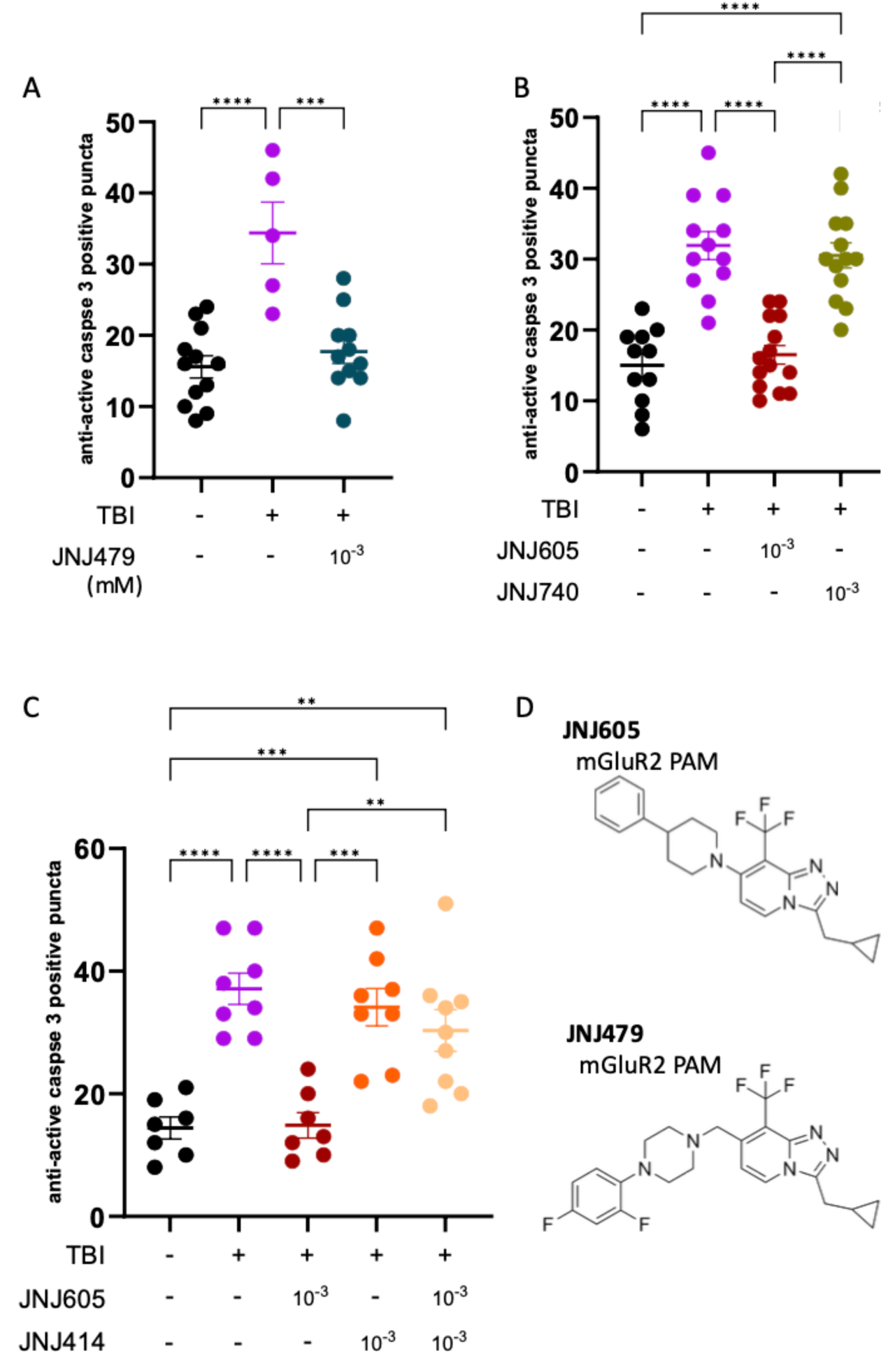
Specificity of JNJ-42153605 action as a mGluR2 PAM is supported by active analogs, inactive compounds at mGlu2 PAM, and opposing actions of an mGluR2 NAM. **A.** A second mGluR2 Positive Allosteric Modulator (PAM) prevents TBI-induced cell death. Applying the mGluR2 PAM JNJ-46356479 (**JNJ479**) following traumatic brain injury (TBI) reduces cell death similar to JNJ605. **B.** An inactive compound at mGluR2, (**JNJ740)**, induces no apparent impact on cell death following TBI. When co-applied with JNJ605, JNJ740 has no apparent effect on the outcomes. **C.** An mGluR2 negative allosteric modulator (NAM), **JNJ414,** does not significantly alter cell death following TBI. This mGluR2 NAM significantly reverses the protective effect of the PAM JNJ605. **D.** The mGluR2 PAMs used in this study have distinct structures. N=8-16 larva per treatment. Each symbol represents an individual animal, mean ±SE plotted. *p<0.05, **p<0.01 ***P<0.001, ****p<0.0001 by Kruskal-Wallis test and Dunn’s multiple comparisons test post-hoc.

Next, we compared the protective effects of JNJ605 versus an inert analog, JNJ740. JNJ605 was significantly more efficacious than JNJ740 in reducing TBI-induced cell death (**Fig 3B**). There was no evidence in our data that the inert analog JNJ740 impacts cell death induced by TBI.

To further consider the specificity of JNJ605 in the zebrafish system, we applied an mGluR2 NAM, JNJ414. Alone, JNJ414 had no apparent impact on TBI-induced cell death at the single dose evaluated (**Fig 3C**). Next, we determined if the actions of JNJ605 and JNJ414 act in opposition. JNJ605 again reduced TBI-induced cell death (as in Fig 2 and 3B), but this action was significantly reduced by the simultaneous application of JNJ414 (**Fig 3C**). This demonstrates that the protective actions of JNJ605 we report occur principally via its known role as an mGluR2 PAM, because any action via other molecular targets would be unlikely to be countered by an mGluR2 NAM.

### Restoring seizures reverses the beneficial effects of LEV and JNJ605

Many intertwined events contribute to the neurodegenerative outcomes following neurotraumatic insult. These at least include the many molecular events of neuroinflammation and hypoxia, which may serve to mutually amplify post-traumatic seizures and excitotoxicity in a feedforward manner (**Fig 4A**). Moreover, following neurotrauma JNJ605 and LEV could potentially impact these and/or other pathomechanisms directly. We confirmed that LEV and JNJ605 are effective in reducing seizures in zebrafish larvae (**Supp Fig S5**). Next, we reasoned that if JNJ605 and/or LEV are exerting their protective effects on tau aggregation via mitigating post-traumatic seizures, then a reinstatement of seizures in our paradigm would negate the compounds’ therapeutic actions. We applied the convulsant kainate following TBI, and observed significantly exacerbated tau aggregation (**Fig 4B,C**; approximate doubling of tau puncta. p<0.01). Co-application of kainate with LEV reversed the protective effect of LEV on TBI-induced tau aggregation (**Fig 4B**), increasing tau puncta to levels similar to animals treated with TBI and vehicle alone. Similarly, kainate reversed the protective effects of JNJ605 (**Fig 4C**). The data is consistent with our working model, that the neural hyperexcitability induced by TBI links the neurotrauma to later tau aggregation, and this early mechanism is near the top of a hierarchy of other likely pathomechanisms that may also promote tau aggregation (**Fig 4A**).

**Figure 4.**
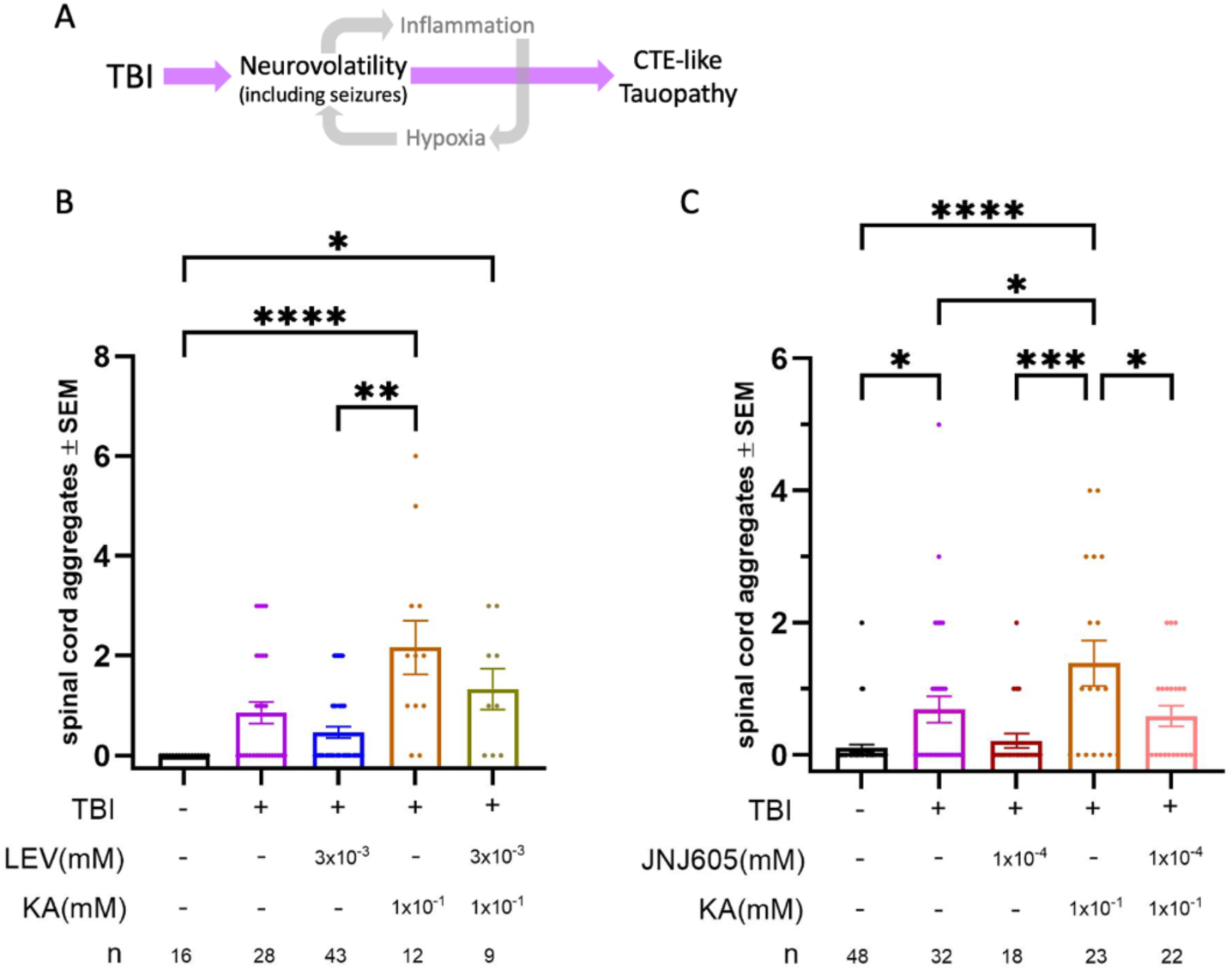
Mitigating Post-traumatic Seizures is a candidate mechanism how Levetiracetam (LEV) and JNJ-42153605 pre-treatments prevent subsequent tau aggregation. **A.** Our model speculates that neural hyperexcitability/ post-traumatic seizures is one of several feed-forward pathomechanisms that amplify one-another and link TBI to CTE-like tau aggregation. If neural hyperexcitability is near the top of this hierarchy, then blocking and restoring it should mitigate and restore tau aggregation, respectively. Our data address events in the days following TBI, and suggest further work is warranted that considers these during disease progression. **B, C.** LEV is typically considered for its actions as an anti-epileptic drug (AED) but its protective impacts on degenerating brains may occur via various mechanisms. The convulsant kainate (KA) was used to sustain seizures in TBI-treated larvae. TBI induced tau aggregation in larvae, and this was significantly exacerbated in larvae receiving KA after TBI. **B.** Effective doses of LEV reduced tau aggregation in a manner that was reversed by co-application of KA, suggesting that LEV is unable to prevent tau aggregation when LEV’s impacts on seizures are muted. **C.** JNJ-42153605 (“JNJ605”, an mGluR2 positive allosteric modulator) reduced tau aggregation in a manner that was reversed by co-application of KA, speculatively suggesting that JNJ605’s protective mechanism is related to mitigating seizure-like events. Each symbol represents an individual animal, n>18 individuals per treatment and data was derived from three separate experiments. Horizontal lines plot the mean ±SE. *p<0.05, **p<0.01, ***P<0.001, ****p<0.0001 by Kruskal-Wallis test and Dunn’s multiple comparisons test post-hoc.

## DISCUSSION

Forms of neural hyperexcitability including post traumatic seizures are emerging as substantial mechanisms that link TBI neurotrauma to subsequent dementias such as CTE and AD[2]. That mechanism is enticing because it is druggable, including via re-evaluating the AEDs used following TBI to consider not only their impacts on PTS, but also their impacts on longer-term outcomes including tauopathy. Here, multiple lines of evidence support that the AED LEV alone, and when combined with mGluR2 PAMs, have promise to prevent tau aggregation. The mGluR2 PAM JNJ605 was on its own able to mitigate tau aggregation following TBI at sub-mM doses. LEV was similarly effective in reducing TBI-induced tau aggregation. The data are derived from 9-point and 11-point dose-response outcomes evaluating hundreds of individual animals, enabled by a genetically encoded tau aggregation biosensor expressed in the CNS neurons of transparent larval zebrafish. The data address the early events following TBI, and suggest that further experiments are warranted in rodent models and with longer time courses that consider the progressive nature of these diseases.

LEV and JNJ605 both act principally at presynaptic targets and dampen synaptic vesicle release. Considering previous preclinical works that demonstrated potent synergism of LEV and mGluR2 PAM in dampening seizures in mouse models of epilepsy[16, 17], we examined whether the protective effects of LEV and JNJ605 would amplify each other in mitigating TBI-induced tau aggregation. Again turning to tau aggregation quantification in hundreds of larval zebrafish, the data show that a low-moderate dose of LEV can substantially reduce the dose of JNJ605 that is required to prevent tau aggregation. The converse was confirmed in an independent dataset, wherein a low dose of JNJ605 was sufficient to reduce the dose of LEV that is sufficient to abrogate tau aggregation following TBI.

### AEDs as dementia prophylactics

AEDs applied following TBI, perhaps for short durations of weeks or in longer-term applications, may offer hope to prophylactically prevent later dementias. R&D has found success in the treatment of stroke, where mitigating the neurotrauma in a timely fashion (e.g. via stroke ambulances) has not only acute benefits in preventing death and disability, but also has long-term impacts in mitigating the subsequent and progressive cognitive decline. Thus, we suggest that research in AEDs applied following TBI may benefit not only the impacts on PTS and PTE, but also mitigate later consequences on cognitive health.

At this time, two AEDs in the literature are candidates to prophylactically mitigate dementia following TBI. These are retigabine (RTG) and LEV [2]. We previously assessed RTG, a K_V_7 channel blocker that acts to reduce rates of action potential firing. In our zebrafish paradigm, RTG effectively mitigated TBI-induced tau aggregation [1]. Independently, RTG was utilized in mouse models of blast TBI. Application of RTG for only days following TBI and was successful in muting several pathological hallmarks of dementia and cognitive decline several months after TBI in mice[3]. This suggests that a conserved and important pathomechanism exists between disparately related vertebrate models. Moreover, precedent exists where pharmacology insights from our larval zebrafish paradigm (with a short timeline) translate well into mammalian models with a more protracted and progressive dementia.

While past works with RTG in zebrafish and mice were compelling, exploring AEDs with synaptic mechanisms of action was warranted. This is due in part on the need to focus on early intervention during the course of dementia progression, coupled with a substantial body of evidence that synaptic dystrophies are among the earliest events in dementia aetiology. At this time it is unclear if AEDs will be useful in slowing progression of many types of dementia, and this may very much depend on the dementia pathomechanism. Further, defining which patients are likely to benefit from AED treatment may be a substantial challenge. We focus our efforts here on preclinical models of TBI because the neurotrauma is well known to instigate various forms of neural hyperexcitability including PTS; this provides a known onset of events and a rationalized timeline of AED application. Thus, we speculated that synapse-targetted AEDs that can blunt PTS immediately after TBI would be worth testing as potential dementia prophylactics[2]. The early success with LEV in studies encouraged us to prioritize its application, and to subsequently explore mechanisms to amplify LEV’s potency so as to minimize its potential harms if applied at higher doses.

LEV has been shown to have good potential in slowing cognitive decline and muting AD-related hippocampal activity in rodent models of dementia[7, 22]. LEV is also promising in clinical trials with an intriguing subset of patients who are at risk of developing AD[14, 23]. The subset of MCI patients who exhibited neural hyperexcitability (detected with overnight EEG) benefited from LEV compared to placebo, whereas MCI patients that did not exhibit such neural hyperexcitability were not shown to suggest a positive impact of LEV[14, 23]. This suggests that LEV may benefit an important subset of patients if applied early in dementia progression, perhaps including the subset of patients with epileptic Alzheimers disease, neural hyperexcitability and perhaps relating to a history of TBI. In the present study, LEV was remarkably successful as a prophylactic treatment following TBI, such that moderate doses were able to block the subsequent onset of tau aggregation and cell death.

Despite LEV being generally well-tolerated, its potential as a dementia prophylactic must be balanced against its potential adverse effects, alongside the likelihood of mitigating any such harms. LEV has been associated with drowsiness, dizziness, fatigue and irritability in some patients[15, 24, 25]. Clinically, AEDs are often applied in combinations and we speculated that a polypharmacy approach may be able to improve LEV’s efficacy, and thus assist in mitigating these concerns.

Here we tested the ability of mGluR2 PAMs to improve LEV’s efficacy. Elegant studies in mouse models of seizure have shown a dramatic ability of mGluR2 PAMs, JNJ605 and JNJ479, to synergize with LEV in its antiepileptic outcomes[4, 16]. PAMs are well suited to such a strategy because they allow orthosteric signalling of glutamate continue, and they are less prone to tachyphylaxis compared to orthosteric drugs. In our study, JNJ605 was successful in mitigating subsequent tau aggregation following TBI. Co-treatment with JNJ605 improved the prophylactic efficacy of LEV by at least an order of magnitude.

### Limitations of the study

The tau aggregation outcome we quantified in our preclinical model is not mainstream, but it is well-validated. The Tau4R-GFP reporter has been informative when expressed in cultured cells. Past work in zebrafish characterized our genetically encoded Tau4R-GFP reporter as being an intact protein, and induced to aggregate by homogenate from mouse brains laden with human tau pathology (but not control brains) or by human tau fibrils (but not monomers)[1]. Regardless, it will be important to validate these outcomes using more mainstream endpoints of tauopathy such as assessing tau biochemistry, silver staining and Tau phosphorylation. Additionally, it is possible that the treatments we applied alter the behaviour of the reporter; we could find no evidence of this at high doses of the drugs (see Methods) and this explanation would need to be true of multiple treatments (while coincidentally mitigating cell death measures) so we deem it unlikely. The tau aggregation reported here is known to vary systematically vs. TBI dose, is induced by delivery of human tau in carefully controlled experiments, and accumulates progressively over time[1, 19, 20].

Moreover, our model provides an ethically favourable approach (using larval zebrafish to replace neurotraumatic insults on adult fish or mammals) that allows robust sample sizes thereby implementing expansive dose-response curves to characterize *in vivo* drug actions. A principal limitation of our approach is the short timeline between TBI and biomarker assessment; we determine TBI outcomes days after neurotrauma. The data may be modelling disease onset moreso than late disease progression which may be emergently more complex. We are encouraged, however, that the one AED previously identified by our approach, RTG[1], has been translated to mouse and improved outcomes months after TBI despite only brief application following injury[3]. Further, LEV shows promise to improve longer-term cognitive outcomes in murine models and in clinical trials, though a paucity of similar works are available in rodents regarding mGluR2 PAMS[26].

The current data do not fully address the concept that seizures are a link between TBI and the outcomes reported, though that is addressed elsewhere as reviewed above. In particular the current work did not report the level or time-course of post-traumatic seizures, and correlating those to tauopathy and cell death outputs would be worthwhile. Thus, it remains possible that the antiepileptic LEV acts through non-canonical mechanisms; these hypothetical mechanisms are, at least in the current model, able to be overcome by the convulsant kainate. Further work addressing the timing and role of seizure-like events following TBI is urgently needed.

### Summary

Our data suggest that AEDs that dampen presynaptic vesicle release can impact the early events in the neurotrauma cascade. Other pathomechanisms are certainly very important, including hypoxia and neuroinflammation that amplify each other and neural hyperexcitability in a vicious cycle (Fig 4A). Speculatively, it appears that neural hyperexcitability may be early in this cycle, and high up in a hierarchy of aetiological events, because blocking post-traumatic seizures is sufficient to prevent late dementias. Moreover, neural hyperexcitability following TBI is a sufficient pathomechanism insomuch that replacing seizures during such treatments leads to the reappearance of later tauopathy (Fig 4).

Future work is warranted on LEV and mGluR2 PAMs as potential prophylactics, especially in situations with prominent neural hyperexcitability such as tau aggregation that develop following TBI. This future work must include assessment in (rodent) paradigms where long-term administration can be monitored for safety and efficacy.

## MATERIALS and METHODS

### Zebrafish Husbandry

All procedures for care of zebrafish were approved by the University of Alberta via their Animal Care and Use Committee:BioSciences that operates within the guidelines and oversight of the Canadian Council on Animal Care. Transgenic zebrafish were maintained in the Casper background, expressing our previously described [1] genetically-encoded fluorescent reporters in the CNS neurons. Tauopathy reporter Tau4R-GFP is a fusion of the human Tau repeat region [*Tg(eno2:Hsa.MAPT_Q244-E372−EGFP)ua3171* larvae (ZFIN ID: ZDB-ALT-211005-6)], and is co-expressed with full-length human tau (no GFP physically linked; *Tg(eno2:hsa.MAPT-ires-egfp)Pt406* larvae (ZFIN ID: ZDB-ALT-080122–6).

### Materials

LEV was purchased from Millipore Sigma, product L8668 at ≥98% purity. JNJ605 was purchased from MedChemExpress, product HY-18162 at >98% purity. An alternative preparation of JNJ605 dubbed “JNJ340” in Supplemental Figure S3 was provided as a blinded sample from Janssen Research & Development division, along with blinded samples of inactive compounds, JNJ414 and JNJ472.

*Quantifying tau aggregation and compound application*Zebrafish larvae at 3 days post fertilization (dpf) expressing the tau biosensor were subjected to our TBI assay (10mL syringe, foam block holder, 5x drop of 300g weight, 48” height) that is described in two publications devoted to methodological details and three publications that validate the approach in multifaceted ways [1, 19, 20]. The approach loads larval zebrafish, in their typical bath media, into a syringe. A controlled weight is dropped on the syringe and delivers a pressure wave through the animal and thus is likely to damage all organs and tissues akin to a blast injury. Animals are repositioned in the syringe between each of five weight drops; this strategy reduces the position of animals as a confound between individuals. This injury dose has minimal impacts on survival while inducing substantial seizure-like and pathological outcomes. Other workers have confirmed this approach leads to behavioural deficits and seizure-like hyper-activity [27].

Larvae were treated with LEV, JNJ605, or a combination of LEV + JNJ605 in the concentrations indicated for 38-44 hours, beginning 30-90 minutes post injury. Drug dilutions were made from 300 mM LEV (miliQ solvent) and 80 mM JNJ605 (DSMO solvent) stock stored in −80 C. Stock solutions were diluted in 5 mL of normal fish water (E3 media) to 2x the final concentrations, which was then added to 5mL E3 media in 60×15 mm petri dishes containing the larvae. DSMO was added to each group to match the highest DSMO concentration from JNJ605 dilutions (0.0125% DSMO.) Bath solutions were not changed until 38-44 hours before resuming regular daily media changes. Petri dishes held a maximum of 30 larvae. Larvae were randomly placed into petri dish treatment group equally following each TBI event. Fluorescent puncta of Tau-GFP were quantified, by observers who were blinded to the treatments (TBI and drugs), in the larval spinal cord using a fluorescent microscope (Leica M165 FC). GFP-positive puncta in the spinal cord were counted, anterior of the hindbrain and through to the posterior tip. Puncta with autofluorescence were excluded. Larvae with notable morphological defects prior to TBI were excluded. We did not note any changes to animal survival, morphology, or developmental rate, including at the high doses applied that were several orders of magnitude more concentrated than the efficacious doses.

To test if the reduced tau puncta during treatments applied reduced the abundance of the Tau-GFP reporter in the CNS, we applied relatively high doses (LEV at 3×10^-4^ mM, JNJ605 at 10^-5^ mM). We collected images of the spinal cord using microscopy setting equivalent to our screening, and quantified GFP fluorescent intensity using ImageJ. Spinal cords receiving vehicle vs. drugs showed mean fluorescence of 1.1±0.14 vs. 1.5±0.12, respectively (arbitrary units, mean±SD, n=9 or 10 fish/group). In sum, we could find no evidence that the compounds applied decreased the Tau4R-GFP fluorescent intensity.

### Immunohistochemistry

Cell death was detected using immunohistochemistry. Zebrafish larvae at 3dpf underwent TBI as above and were treated with any indicated drugs for 44 hours. After drug removal at 5dpf larvae were euthanized and fixed in 4% PFA overnight. Subsequently, the larvae were permeabilized and washed with primary and secondary antibodies over two days. Larvae received primary antibody anti-active-caspase-3 at 1:500 dilution (raised in rabbit, BD Pharmingen # 559565) and the secondary antibody anti-rabbit AlexaFluor488 (raised in goat, Invitrogen # A32731).

### Statistics

All statistical analyses were performed using GraphPad Prism Software (Version 10, GraphPad, San Diego, CA). Experiments were replicated at least twice, with individual larvae being the sampling unit. No outliers or other data were excluded. The experimenters were blinded to the treatments prior to quantifying outcomes. Dose response curves were created using three parameter inhibitor vs. response nonlinear Poisson regression, followed by a likelihood ratio test for comparison of two curves and Goodness-of-fit using Log-likelihood (Nagelkerke’s pseudo-R-squared). For comparison between three or more groups, ordinary one-way ANOVA followed by post-hoc Tukey’s multiple comparisons test or Kruskal-Wallis test followed by post-hoc Dunn’s multiple comparison tests (for nonparametric data) were performed. For analysis of binary data, multiple logistic regression was performed with area under the ROC curve, Log-likelihood (G-squared), and Nagelkerke’s R-squared (Pseudo R-squared) calculations to assess model fit.

## ACKNOWLEDGEMENTS

We are grateful to Lance Doucette, Wil Kinley and Tanja Zerulla for technical assistance and early discussions on the work. Science Animal Support Services at the University of Alberta were instrumental to success of the work.

---

**Figure S1.**
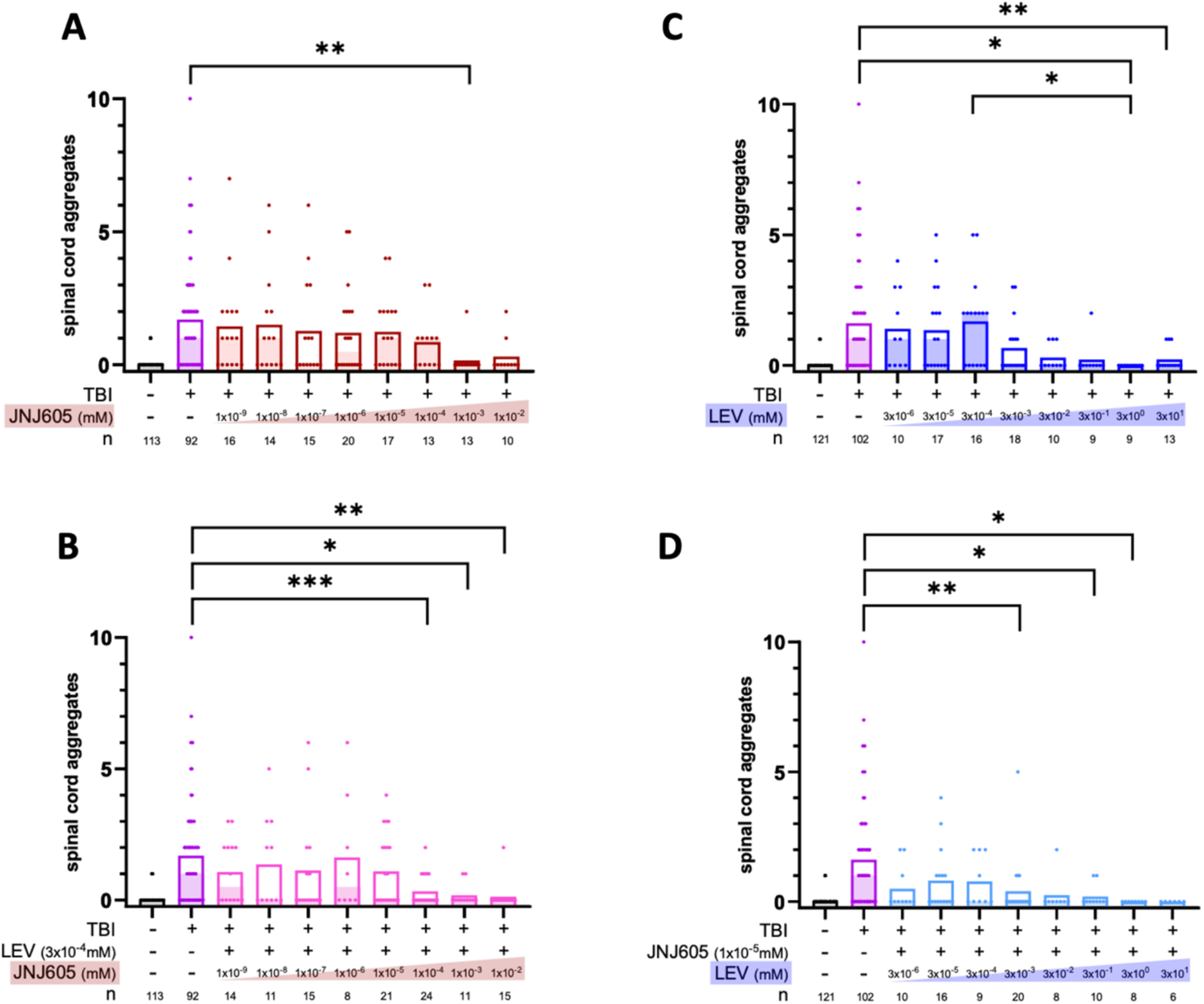
Raw data used to generate. Figure 1**. JNJ605 and LEV applied after TBI potentiate reduction in injury-induced tau aggregation.** Each dot represents the abundance of Tau puncta in an individual larvae. Empty bars in each data set represent the mean (variance around the mean is in Fig 1), whereas the shaded bar represents the median. Sample sizes (n values) at the bottom of the graphs are the number of individual animals quantified for each treatment dose; most doses were assessed on multiple days and data was combined. The larvae analyzed here came from multiple independent clutches of transgenic fish analyzed on many separate days, including 7 clutches for LEV alone (data in dark blue); 5 clutches for LEV when added to a constant dose of JNJ605 (light blue); 8 clutches for JNJ605 alone (red); 5 clutches for when JNJ605 was added to a constant dose of LEV (pink). For easy reference, baseline data (without drug addition) is replicated on panels A and B, and on panels C and D. *p<0.05, **p<0.01, ***P<0.001 by Kruskal-Wallis test with Dunn’s multiple comparisons post-hoc test.

**Figure S2.**
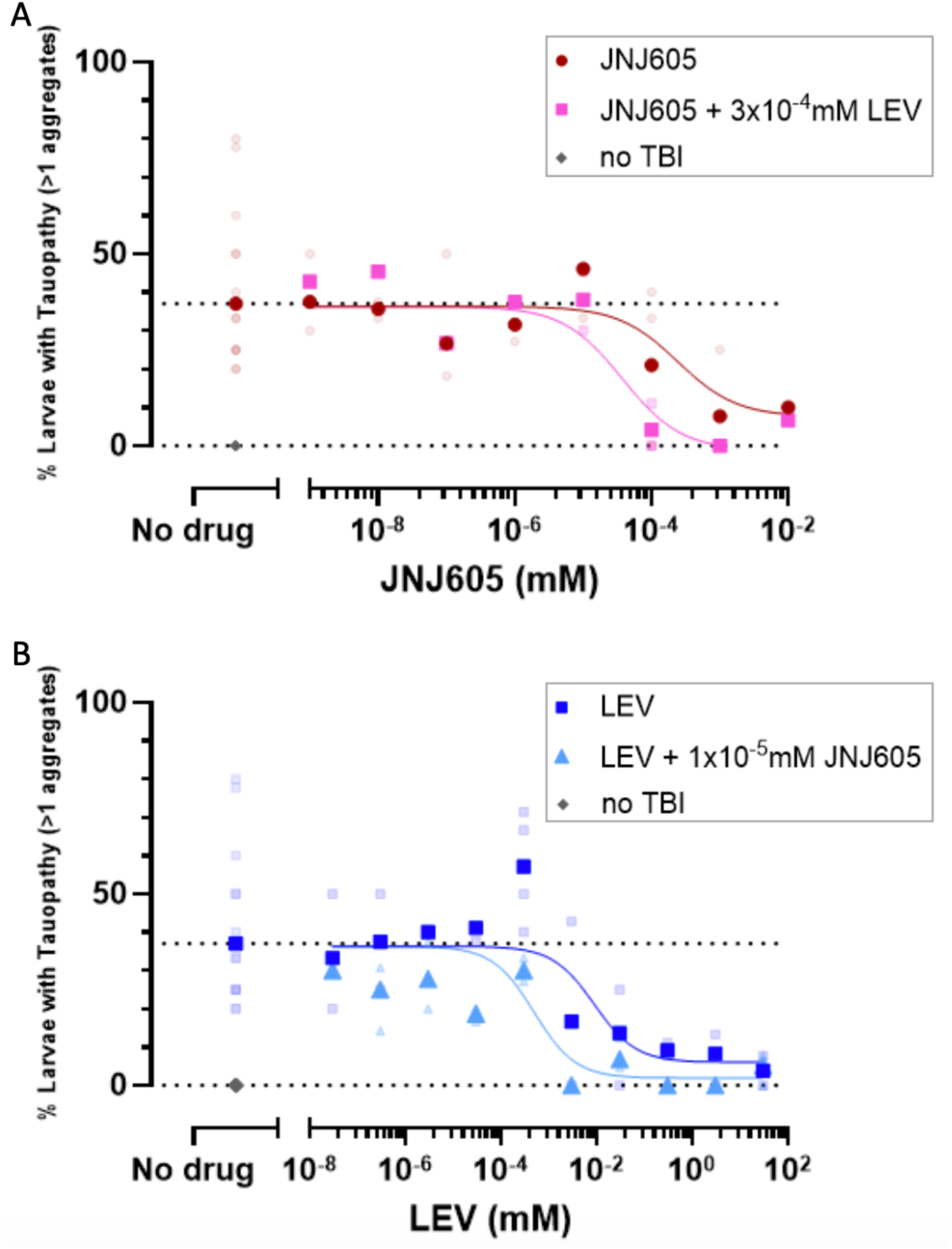
Levetiracetam and the mGluR2 PAM JNJ-42153605 applied following TBI interact to mitigate subsequent tau aggregation. Separate analysis of the same fish reported in Figure 1 and S1, but re-analyzed to consider the <u>percentage of animals</u> showing elevated tau puncta in each trial. Tau levels were considered elevated if >1 tau puncta was observed in the animal. Statistical analysis of this data considered the odds ratio (OR) varying with drug concentration. **A.** JNJ-42153605 (JNJ605) reduces the proportion of fish with tau aggregation in a dose-dependent manner, and a low dose of Levetiracetam (LEV) significantly potentiates this effect. JNJ605 increased the odds of larvae being tau free post-TBI 1.2-fold/µM (OR 1.218, 95% CI 1.008-1.746). Co-treatment with subeffective 3×10^-4^ mM LEV appeared to increase the odds of larvae being tau free post-TBI about 1.4-fold compared to larvae without LEV co-treatment but the result was non-significant with a lower confidence interval <1 (OR 1.443, 95% CI 0.8702-2.443). **B.** LEV reduces the proportion of fish with tau aggregation in a dose-dependent manner, and a low dose of JNJ605 significantly potentiates this effect. LEV increased the odds of larvae being tau free post-TBI by 1.11-fold/mM (OR 1.11, 95% CI 1.041-1.271). Co-treatment with subeffective 10^-5^ mM JNJ605 increased the odds of larvae being tau free post-TBI by 2.694-fold compared to larvae without JNJ605 co-treatment (OR 2.694, 95% CI 1.674-4.459). Each dark symbol is the mean across independent trials on separate days; each lighter symbols is the raw data from a single trial. Animals (larvae) with >1 tau aggregate in the CNS were included. For easy reference, baseline data (without drug addition) is replicated on panels A and B. Line of best fit curve generated by three parameter inhibitor vs response nonlinear regression. Statistical analysis was performed using a multiple logistic regression (A. AUC = 0.6134, P< 0.001. G-squared = 12.55, P<0.01, Nagelkerke’s pseudo R-squared 0.04587; B. AUC = 0.7057, P<0.001. G-squared = 33.83, P <0.0001, Nagelkerke’s pseudo R-squared 0.09687).

**Figure S3.**
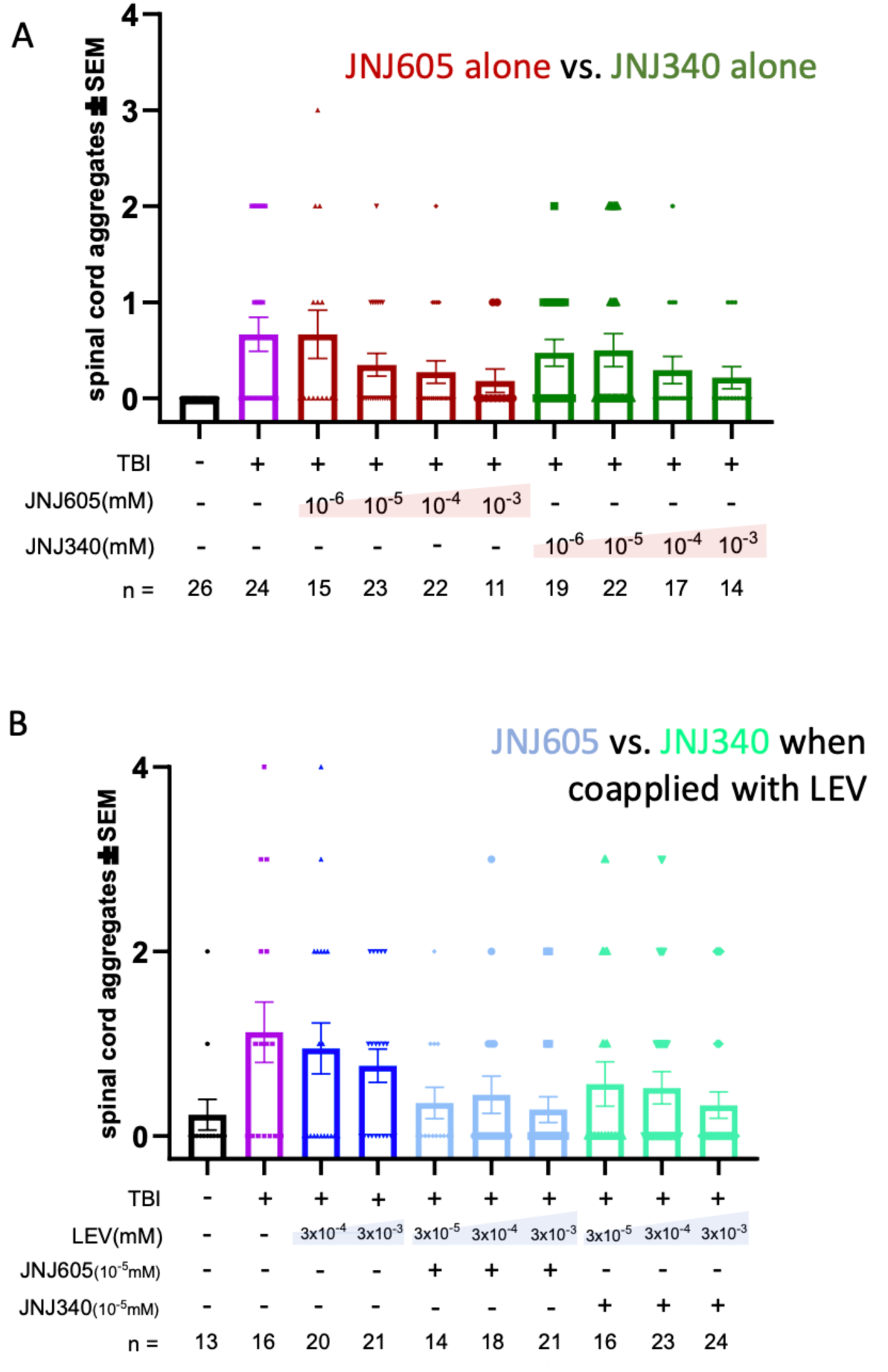
Independent sources of mGluR2 PAM JNJ-42153605 are both efficacious when applied following TBI interact to mitigate subsequent tau aggregation. This data reports two independent replications separate from the data in experiment in Figure 1, showing JNJ605 efficacy is consistent regardless of its source. Commercially available JNJ-42153605 from MCE is labelled as “JNJ605” (used throughout manuscript). JNJ-42153605 supplied by Janssen was denoted here as “JNJ340” for the purpose of this experiment. Impacts of JNJ605 on mitigating tau puncta and on potentiating LEV’s efficacy is consistent between sources. Compound’s efficacy is not significantly different when comparing between the two compound sources (Kruskal-Wallis test, Dunn’s multiple comparisons test post-hoc). Each dot represents the abundance of Tau puncta in an individual larvae. Bars represent mean ±SE.

**Figure S4.**
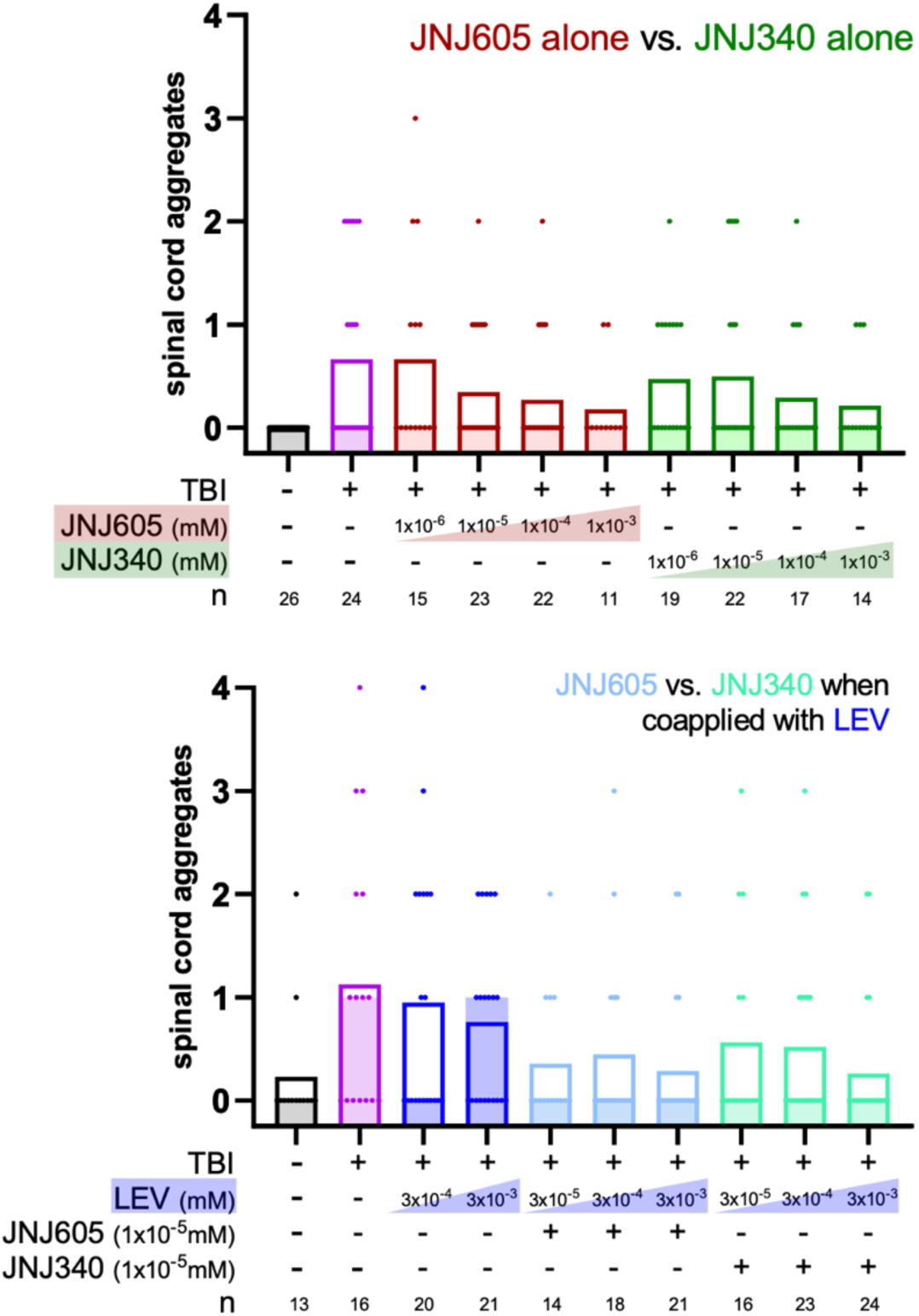
Independent sources of mGluR2 PAM JNJ-42153605 are both efficacious when applied following TBI interact to mitigate subsequent tau aggregation. The same data as Figure S2 replotted to report the median (shaded bars). Commercially available JNJ-42153605 from MCE is labelled as “JNJ605” (used throughout manuscript). JNJ-42153605 supplied by Janssen was denoted here as “JNJ340” for the purpose of this experiment. Impacts of JNJ605 on mitigating tau puncta and on potentiating LEV’s efficacy is consistent between sources. Compound’s efficacy is not significantly different when comparing between the two compound sources (Kruskal-Wallis test, Dunn’s multiple comparisons test post-hoc). Each dot represents the abundance of Tau puncta in an individual larvae. Bars represent the mean (variance is reported in Fig S3), and shaded bars represent the median.

## REFERENCES

1. Alyenbaawi, H., et al., Seizures are a druggable mechanistic link between TBI and subsequent tauopathy. Elife, 2021. 10.

2. Locskai, L.F., H. Alyenbaawi, and W.T. Allison, Antiepileptic Drugs as Potential Dementia Prophylactics Following Traumatic Brain Injury. Annu Rev Pharmacol Toxicol, 2024. 64: p. 577–598.

3. Vigil, F.A., et al., Acute Treatment with the M-Channel (Kv7, KCNQ) Opener Retigabine Reduces the Long-Term Effects of Repetitive Blast Traumatic Brain Injuries. Neurotherapeutics, 2023. 20(3): p. 853–869.

4. Zhang, Y., et al., Connectivity Mapping Using a Novel sv2a Loss-of-Function Zebrafish Epilepsy Model as a Powerful Strategy for Anti-epileptic Drug Discovery. Front Mol Neurosci, 2022. 15: p. 881933.

5. Atwood, R., et al., Use of Levetiracetam for Post-Traumatic Seizure Prophylaxis in Combat-Related Traumatic Brain Injury. Mil Med, 2023. 188(11-12): p. e3570–e3574.

6. Browning, M., et al., Levetiracetam Treatment in Traumatic Brain Injury: Operation Brain Trauma Therapy. J Neurotrauma, 2016. 33(6): p. 581–94.

7. Caudle, K.L., et al., Neuroprotection and anti-seizure effects of levetiracetam in a rat model of penetrating ballistic-like brain injury. Restor Neurol Neurosci, 2016. 34(2): p. 257–70.

8. Chen, Y.H., et al., Levetiracetam prophylaxis ameliorates seizure epileptogenesis after fluid percussion injury. Brain Res, 2016. 1642: p. 581–589.

9. Javed, G., et al., Use of Levetiracetam in Prophylaxis of Early Post-Traumatic Seizures. Turk Neurosurg, 2016. 26(5): p. 732–5.

10. Jones, K.E., et al., Levetiracetam versus phenytoin for seizure prophylaxis in severe traumatic brain injury. Neurosurg Focus, 2008. 25(4): p. E3.

11. Nita, D.A. and C.D. Hahn, Levetiracetam for Pediatric Posttraumatic Seizure Prophylaxis. Pediatr Neurol Briefs, 2016. 30(3): p. 18.

12. Pearl, P.L., et al., Results of phase II levetiracetam trial following acute head injury in children at risk for posttraumatic epilepsy. Epilepsia, 2013. 54(9): p. e135–7.

13. Zampella, B., et al., Seizure Prophylaxis in Traumatic Brain Injury: A Comparative Study of Levetiracetam and Phenytoin Cerebrospinal Fluid Levels in Trauma Patients with Signs of Increased Intracranial Pressure Requiring Ventriculostomy. Cureus, 2019. 11(9): p. e5784.

14. Vossel, K., et al., Effect of Levetiracetam on Cognition in Patients With Alzheimer Disease With and Without Epileptiform Activity: A Randomized Clinical Trial. JAMA Neurol, 2021. 78(11): p. 1345–1354.

15. Lin, C.Y., M.C. Chang, and H.J. Jhou, Effect of Levetiracetam on Cognition: A Systematic Review and Meta-analysis of Double-Blind Randomized Placebo-Controlled Trials. CNS Drugs, 2024. 38(1): p. 1–14.

16. Metcalf, C.S., et al., Potent and selective pharmacodynamic synergy between the metabotropic glutamate receptor subtype 2-positive allosteric modulator JNJ-46356479 and levetiracetam in the mouse 6-Hz (44-mA) model. Epilepsia, 2018. 59(3): p. 724–735.

17. Metcalf, C.S., et al., Efficacy of mGlu(2) - positive allosteric modulators alone and in combination with levetiracetam in the mouse 6 Hz model of psychomotor seizures. Epilepsia, 2017. 58(3): p. 484–493.

18. Mani, V. and S. Rashed Almutairi, Impact of levetiracetam on cognitive impairment, neuroinflammation, oxidative stress, and neuronal apoptosis caused by lipopolysaccharides in rats. Saudi Pharm J, 2023. 31(9): p. 101728.

19. Gill, T., et al., Delivering Traumatic Brain Injury to Larval Zebrafish. Methods Mol Biol, 2024. 2707: p. 3–22.

20. Locskai, L.F., et al., A larval zebrafish model of traumatic brain injury: optimizing the dose of neurotrauma for discovery of treatments and aetiology. Biol Open, 2025. 14(2).

21. Cid, J.M., et al., Discovery of 8-Trifluoromethyl-3-cyclopropylmethyl-7-[(4-(2,4-difluorophenyl)-1-piperazinyl)methyl]-1,2,4-triazolo[4,3-a]pyridine (JNJ-46356479), a Selective and Orally Bioavailable mGlu2 Receptor Positive Allosteric Modulator (PAM). J Med Chem, 2016. 59(18): p. 8495–507.

22. Isla, A.G., et al., Low dose of levetiracetam counteracts amyloid beta-induced alterations of hippocampal gamma oscillations by restoring fast-spiking interneuron activity. Exp Neurol, 2023. 369: p. 114545.

23. Shandilya, M.C.V., et al., High-frequency oscillations in epileptic and non-epileptic Alzheimer’s disease patients and the differential effect of levetiracetam on the oscillations. Brain Commun, 2025. 7(1): p. fcaf041.

24. Dinkelacker, V., et al., Aggressive behavior of epilepsy patients in the course of levetiracetam add-on therapy: report of 33 mild to severe cases. Epilepsy Behav, 2003. 4(5): p. 537–47.

25. Abou-Khalil, B., Levetiracetam in the treatment of epilepsy. Neuropsychiatr Dis Treat, 2008. 4(3): p. 507–23.

26. Perez-Garcia, G., et al., BCI-838, an orally active mGluR2/3 receptor antagonist pro-drug, rescues learning behavior deficits in the PS19 MAPT. Neurosci Lett, 2023. 797: p. 137080.

27. Wang, K., P. Zhang, and Y. Geng, High-Precision Pneumatic Induction of Traumatic Brain Injury in Larval Zebrafish. Zebrafish, 2026. 23(4-5): p. 132–136.

